# Efficient Visual Perception of Human-Robot Walking Environments using Semi-Supervised Learning

**DOI:** 10.1101/2023.06.28.546903

**Authors:** Dmytro Kuzmenko, Oleksii Tsepa, Andrew Garrett Kurbis, Alex Mihailidis, Brokoslaw Laschowski

## Abstract

Convolutional neural networks trained using supervised learning can improve visual perception for human-robot walking. These advances have been possible due to large-scale datasets like ExoNet and StairNet - the largest open-source image datasets of real-world walking environments. However, these datasets require vast amounts of manually annotated data, the development of which is time consuming and labor intensive. Here we present a novel semi-supervised learning system (ExoNet-SSL) that uses over 1.2 million unlabelled images from ExoNet to improve training efficiency. We developed a deep learning model based on mobile vision transformers and trained the model using semi-supervised learning for image classification. Compared to standard supervised learning (98.4%), our ExoNet-SSL system was able to maintain high prediction accuracy (98.8%) when tested on previously unseen environments, while requiring 35% fewer labelled images during training. These results show that semi-supervised learning can improve training efficiency by leveraging large amounts of unlabelled data and minimize the size requirements for manually annotated images. Future research will focus on model deployment for onboard real-time inference and control of human-robot walking.

## I. Introduction

Computer vision can be used for environment-adaptive control and planning of human-robot locomotion. As an example, previous works [1] have explored the use of smart glasses and wearable inertial sensors to forward predict the leg kinematics during walking for continuous control of robotic prostheses and exoskeletons. Accurate and real-time detection of stairs is especially important for user safety. Vision-based stair recognition systems have used hand-designed feature extractors [2]-[6] and more advanced automated feature engineering via convolutional neural networks (CNNs) [7]-[15] to improve the system design and performance.

Recent studies using deep learning greatly expand on the number of labelled images, most notably *ExoNet* [16] -the largest dataset of egocentric (first-person) images of real-world walking environments. However, the dataset has many environmental classes that overlap with one another with varying degrees, resulting in poor generalization. Kurbis and colleagues [8]-[9] recently developed a 4-class image dataset called *StairNet* based on *ExoNet* by combining similar classes and focusing on stair recognition. The system allowed for more consistent and robust training of a convolutional neural network. Since this system is considered state-of-the-art in vision-based stair recognition and operates on a large-scale open-source dataset, we used their results as a benchmark reference for comparison.

A significant bottleneck of these past studies using images from wearable cameras has been the acquisition of large-scale annotated training data. Manually labelling hundreds of thousands of images is time-consuming and labor intensive. There is a practical need to exploit the unlabelled portions of these image datasets. There have been few attempts to introduce unsupervised learning for image classification of walking environments. For example, [17] used simulated data generation and unsupervised domain adaptation, and [18] generated pseudo-labels to diversify data for different applications, including kinematic gait cycle mode prediction. However, these studies used relatively limited evaluation methods (i.e., small datasets), which can impact their real-world performance and generalization to more complex environments.

Given that *StairNet* includes only a small proportion of the *ExoNet* images (∼10%), we explored the use of the remaining unlabelled data to improve model performance and training efficiency via semi-supervised learning – an advanced method that uses fewer labelled images and large amounts of unlabelled data. Our goal is to support the development of new computer vision systems for environment-adaptive control of robotic prosthetic legs and exoskeletons. This work focuses on improving training efficiency and making computer vision systems more accessible to researchers in wearable robotics by minimizing the number of required labelled images while maintaining high prediction accuracy similar to the previous state-of-the-art [8]-[9].

## II. Methods

### A. Computer Vision Dataset

We used the labelled computer vision dataset called *StairNet* [8]-[9], a derivative dataset of *ExoNet* [16] that focuses on walking environments during stair ascent. The dataset was developed using re-annotated images from six of the twelve original *ExoNet* classes which focus on stair ascent, with more precise cut off points between classes to reduce overlap. The *StairNet* dataset contains ∼515,000 RGB images of four classes, including level-ground (LG), level-ground transition to incline stairs (LG-IS), incline stairs (IS), and incline stairs transition to level-ground (IS-LG). The class distribution of *StairNet* is unbalanced such that the steady-state classes, LG and IS, comprise over 95% of the images, whereas the transition classes, IS-LG and LG-IS, form the remaining 5%.

Our research focuses on using the unlabelled images from *ExoNet* that were not included in the *StairNet* dataset. These images contain walking environments, however, many have obstructions or environments that are not related to stair recognition, as encountered during the *StairNet* development [8]-[9]. These are some of the challenges with using unlabelled images, as prior information of the class distributions or viability of the images is unknown. Note that the images of stairs and level-ground contain real-world environmental features, including other people walking.

### B. Semi-Supervised Learning

We decided to use the FixMatch semi-supervised learning algorithm [19] as a proof-of-concept since it is relatively intuitive and more straightforward to implement compared to more complex algorithms such as self-training with noisy student [20], meta pseudo-labels [21], AdaMatch [22], or contrastive learning for visual representations [23].

Our pipeline is summarized as follows (Figure 1): (1) loading raw labelled and unlabelled images and oversampling with augmentations to the labelled dataset to mitigate false positives during training. (2) Filter and preprocess the unlabelled dataset by retrieving unlabelled image logits via supervised pretrained model and selecting the most probable pseudo-labels that surpass the cutoff parameter *τ*. We preprocess the unlabelled images by applying weak augmentations (i.e., horizontal flips) and strong augmentations (i.e., color intensity, saturation, small rotations and translations, and horizontal flips). We de-fine the batch size ratio parameter *μ* that indicates the difference between the labelled and unlabelled batch sizes. The unlabelled data requires a greater batch size than the labelled dataset such that the bigger the difference, the more impact semisupervised learning can have. We input the labelled and unlabelled batches during training, infer the model on the weakly augmented images, and threshold the received logits by the pseudo-label cutoff parameter *τ*. (3) We train the model by calculating and adding two loss terms: supervised loss (i.e., cross-entropy loss – CE loss) and unsupervised loss (i.e., cross-entropy loss of the thresholded pseudo-label logits calculated against strongly augmented images). The unsupervised loss is scaled by the *λ* parameter that indicates the importance of the unsupervised portion. These semi-supervised parameters (*τ, λ*, and *μ*) are tuned and provide a high degree of model flexibility.

**Figure 1.**
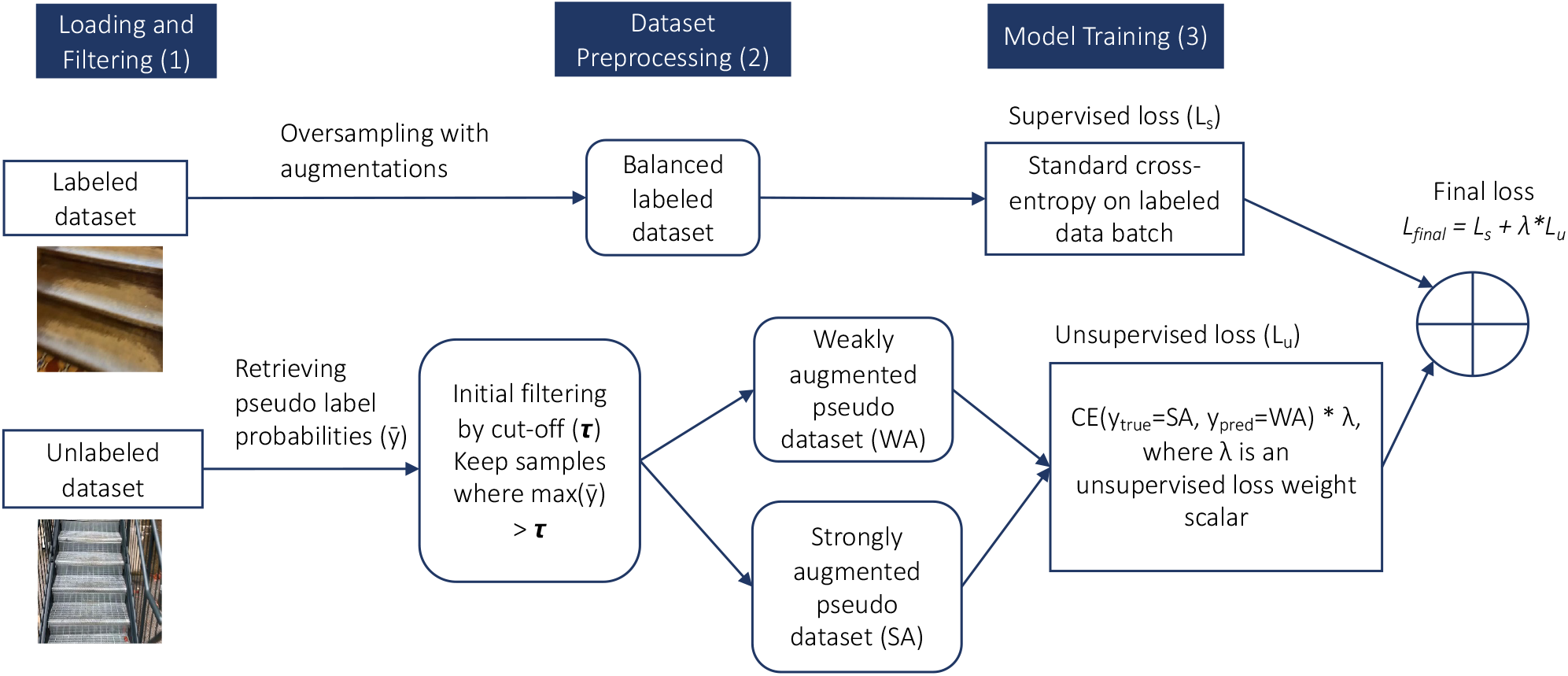
Our semi-supervised learning algorithm with 3 stages: (1) preparing labelled and unlabelled data by oversampling and filtering the non-informative unlabelled images; (2) preprocessing the unlabelled dataset and creating weakly and strongly augmented variations; and (3) using two losses (supervised and unsupervised) to consolidate training and introduce semi-supervised learning.

### C. Mobile Vision Transformer

We developed a vision transformer (ViT) model using the base model of MobileViT [24], which makes use of automated feature engineering similar to standard CNNs. Given that [25] showed that automated deep learning extractors are superior to hand-crafted feature extractors, especially on large-scale image datasets, we focused on convolution and transformer-based models. MobileViT is a transformer-based deep learning model that uses mechanisms of attention and depth-wise dilated convolutions. MobileViT blocks resemble convolutions. The model has low-level efficient convolution and transformer blocks, which can permit MobileViT to run on mobile and embedded computing devices. Its architecture can allow for performance and speed similar to lightweight CNN models like MobileNetV2. We did experiment with MobileNetV3, but found its architecture was not as suitable for our computer vision task and dataset comparable to MobileNetV2.

Three model backbones were considered – MobileViT of sizes XXS, XS, and S, which differ in the number of transformer layers, more sophisticated feature extraction, and pa-rameter count. MobileViT was chosen for its efficient and lightweight design and as a first step toward using transformer models for our computer vision application. Our final transformer-based model was created in TensorFlow 2.0. The model variants were efficiently trained using the high performance computing power of a Google Cloud Tensor Processing Unit (TPU).

### D. Model Training and Optimization

We used the same *StairNet* dataset splits for validation (3.5%) and testing (7%) as Kurbis and colleagues [8]-[9] for benchmark comparison. The labelled training set was initially downsampled to 200,000 images from 461,328 images to allow for greater improvements through semi-supervised learning and to study the impact of reduced labelled data. To address the issue of unknown class distributions and image quality of the unlabelled data, the supervised MobileNetV2 model from [8]-[9] was recreated and trained (Figure 2). This pretrained model was used to retrieve logits of the 4.5 million unlabelled images from *ExoNet*, which were then thresholded in a FixMatch manner. From the unlabelled dataset, ∼1.2 million images surpassed the 0.9 *τ* cutoff. The resulting subset, called *ExoNet-SSL*, had pseudo-labels that closely resembled the original *StairNet* distribution [8]-[9]: IS with 73,404 images (5.5%), IS-LG with 13,640 images (1%), LG with 1,200,017 images (90.1%), and LG-IS with 45,179 images (3.4%) (Table 1).

**Figure 2.**
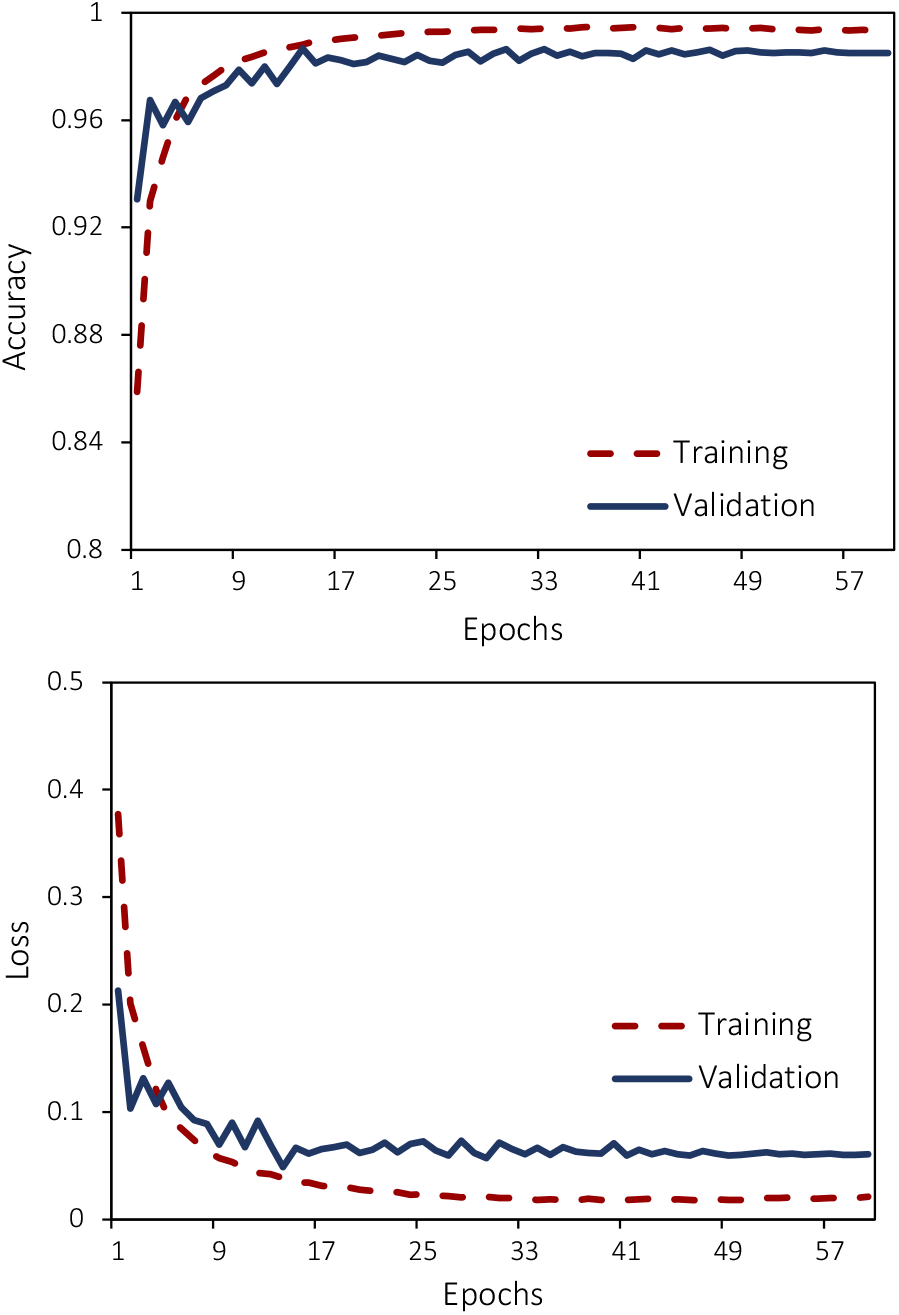
The image classification accuracy and loss for the training and validation sets using the supervised MobileNetV2 model.

**Table 1.**
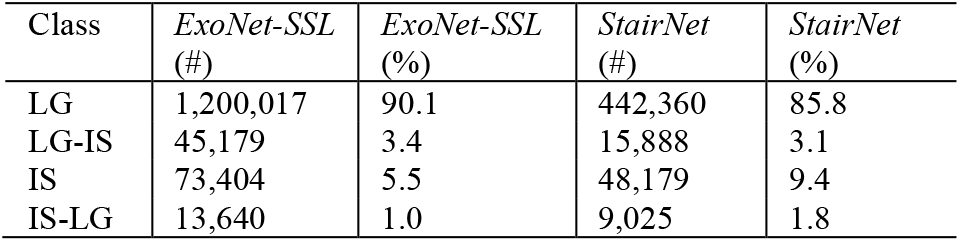
Comparison between the labelled image dataset (*StairNet*) and our new unlabelled dataset (*ExoNet-SSL*) for stair recognition in terms of number of images (#) and percent of the dataset (%).

We experimented with three variants of MobileViT. The lightweight XXS model (∼900,000 parameters) had superior training and inference time but achieved lower categorical accuracy across all classes (accuracy of 97.7% and f1-score of 97.8%). The largest S model (∼4.9 million parameters) trained longer and had slightly weaker performance than XS model (∼1.9 million parameters), which may be explained by overfitting (accuracy of 97.2% and f1-score of 97.3%). Our best model consisted of an ImageNet-pretrained model of MobileViT XS scaled to match our input image resolution of 224×224; stochastic gradient descent (SGD) with 0.9 momentum and Nesterov acceleration; randomly initialized weights; pseudo-label cutoff *τ* of 0.98, a batch difference ratio of 4; unsupervised loss weight λ of 1; initial learning rate = 0.05; labelled batch size of 64; and a FixMatch variation of cosine weight decay learning policy.

The data imbalance was handled by changing the cross-entropy loss to a focal loss [26] with class weight penalization γ of 3 to avoid using oversampling or class weights. We also tested an exponential moving average, which showed that averaged parameters can produce significantly better results than final trained values. The resulting model showed good convergence, but the overall image validation accuracy was inferior to previous vanilla cross-entropy loss experiments.

This leads to the phenomenon that we call “class accuracy allocation”. For our computer vision application, some classes may be considered more important than others for safe human-robot locomotion. The level-ground (LG) class is the most common daily walking environment. Although less common, the transition classes (LG-IS and IS-LG) can have greater implications for falls and user safety. Addressing unbalanced data allows us to redistribute class accuracies, transferring performance from major classes to minor classes. However, this comes with an overall performance degradation as the classes are being more evenly balanced with the focal loss. In other words, the accuracies from major classes (IS and LG) were reallocated to minority classes (IS-LG and LG-IS). The resulting accuracies were 95.1% for IS, 95.8% for IS-LG, 97.5% for LG, and 92.2% for LG-IS.

To combat false positives, we diversified the augmentations applied to the labelled training set, which included minor translations, rotations, contrast, and saturation. Variations of L2 parameter loss and decoupled weight decay [27] were tested. The best models did not include weight decay regularization. Defining a high-performance scheduler for semi-supervised learning can be challenging. We experimented with cosine weight decay as in FixMatch [19] and cosine decay with restarts [28]. The former was more resilient and consistent and was thus selected for our final model design. We performed multiple experiments to find the optimal labelled-unlabelled ratio *μ* and unsupervised loss weight λ parameter. To maximize performance, our final model used 300,000 labelled images (∼65% of the original training labelled dataset) and 900,000 unlabelled images. SGD with Nesterov and momentum was used to optimize the deep learning model with a MobileViT XS backbone. The final set of hyperparameters included a learning rate of 0.045; *τ* = 0.9; a supervised batch size of 64; a batch size ratio *μ* of 3; λ = 1.03; and a cosine decay learning rate schedule. Focal loss was replaced with a categorical crossentropy loss to alleviate degradation in performance caused by class balancing. The final model was trained for 42 epochs (Figure 3).

**Figure 3.**
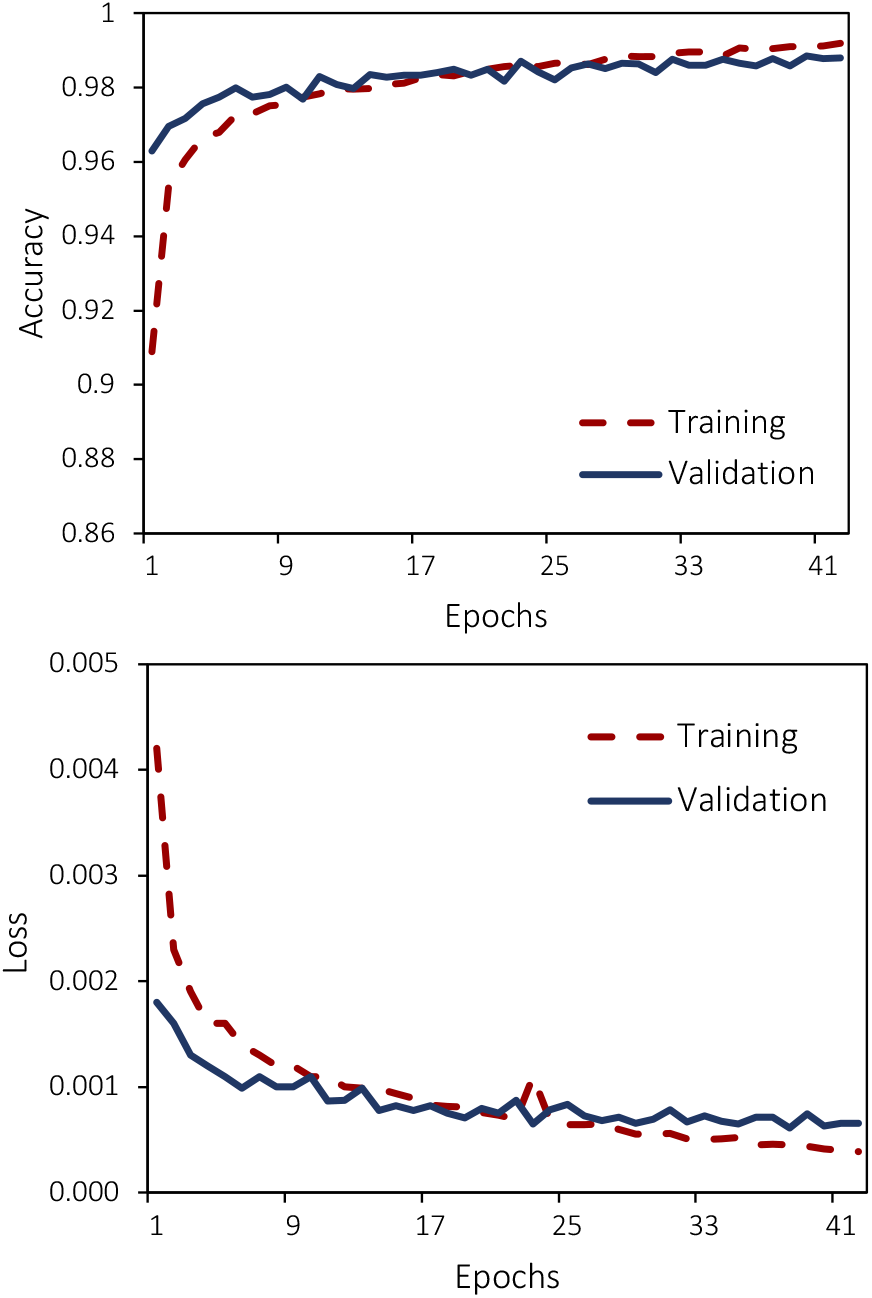
The image classification accuracy and loss for the training and validation sets using our semi-supervised MobileViT model.

## III. Results

The image classification accuracies on the training and validation sets using semi-supervised learning were 99.2% and 98.9%, respectively. When evaluated on the testing set, the model achieved an overall image classification accuracy of 98.8%, a weighted f1-score of 98.9%, a weighted precision value of 98.9%, and a weighted recall value of 98.8%. When compared to standard supervised learning similar to [8]-[9], our semi-supervised learning model achieved 0.4% improvement in overall image classification accuracy and 0.5% improvement in overall weighted f1-score, while requiring ∼35% less manually annotated data (Table 2). Here, classification accuracy is defined as the number of true positives (35,618 images) out of the total number of images in the testing set (36,032 images).

**Table 2.** Comparison between the supervised (MobileNetV2) and semi-supervised (MobileViT XS) learning models in terms of classification performance on the test set and the annotated image requirements. The supervised model results were based on the previous state-of-the-art [8]-[9].

| Training Method | Test Accuracy (%) | f1-score (%) | Precision (%) | Recall (%) | Labelled Images |
| --- | --- | --- | --- | --- | --- |
| Supervised | 98.4 | 98.4 | 98.5 | 98.4 | 461,328 |
| Semi-Supervised | 98.8 | 98.9 | 98.9 | 98.8 | 300,000 |

Tables 3 and 4 show the normalized multiclass confusion matrices for the supervised and semi-supervised learning models, respectively, which show the classification performances on the test set (i.e., new walking environments). For the semisupervised model, the two transition classes (LG-IS and IS-LG) achieved the lowest categorical accuracies (90.6% and 90.4% respectively), which can be attributed to having the smallest class distributions, comprising only 3.1% and 1.8% of the total number of images. Compared to supervised learning, our semi-supervised model achieved higher prediction accuracy for the most highly represented class (LG) while maintaining similar performances on the other walking environments. Figure 4 shows several examples of failure cases whereby our deep learning model misclassified the environment. Nevertheless, it is important to reiterate that our main goal was not necessarily to improve the classification accuracy of individual classes but rather maintain performance while reducing the size of the required labelled training dataset.

**Table 3.** Normalized confusion matrix for the environment predictions (% accuracy) on the test set using the supervised MobileNetV2 model. The horizontal and vertical axes are the predicted and labelled classes, respectively.

|  |  |  |  |  |
| --- | --- | --- | --- | --- |
| IS | 96.9 | 1.5 | 0.3 | 1.3 |
| IS-LG | 6.5 | 90.5 | 2.4 | 0.6 |
| LG | 0.1 | 0.3 | 99 | 0.7 |
| LG-IS | 4.3 | 0.5 | 3.4 | 91.7 |
|  | IS | IS-LG | LG | LG-IS |

**Table 4.** Normalized confusion matrix for the environment predictions (% accuracy) on the test set using the semi-supervised MobileViT model. The horizontal and vertical axes are the predicted and labelled classes, respectively.

|  |  |  |  |  |
| --- | --- | --- | --- | --- |
| IS | 96.9 | 1.1 | 0.5 | 1.5 |
| IS-LG | 5.2 | 90.4 | 3.9 | 0.5 |
| LG | 0.1 | 0.1 | 99.5 | 0.3 |
| LG-IS | 3.4 | 0.4 | 5.6 | 90.6 |
|  | IS | IS-LG | LG | LG-IS |

**Figure 4.**
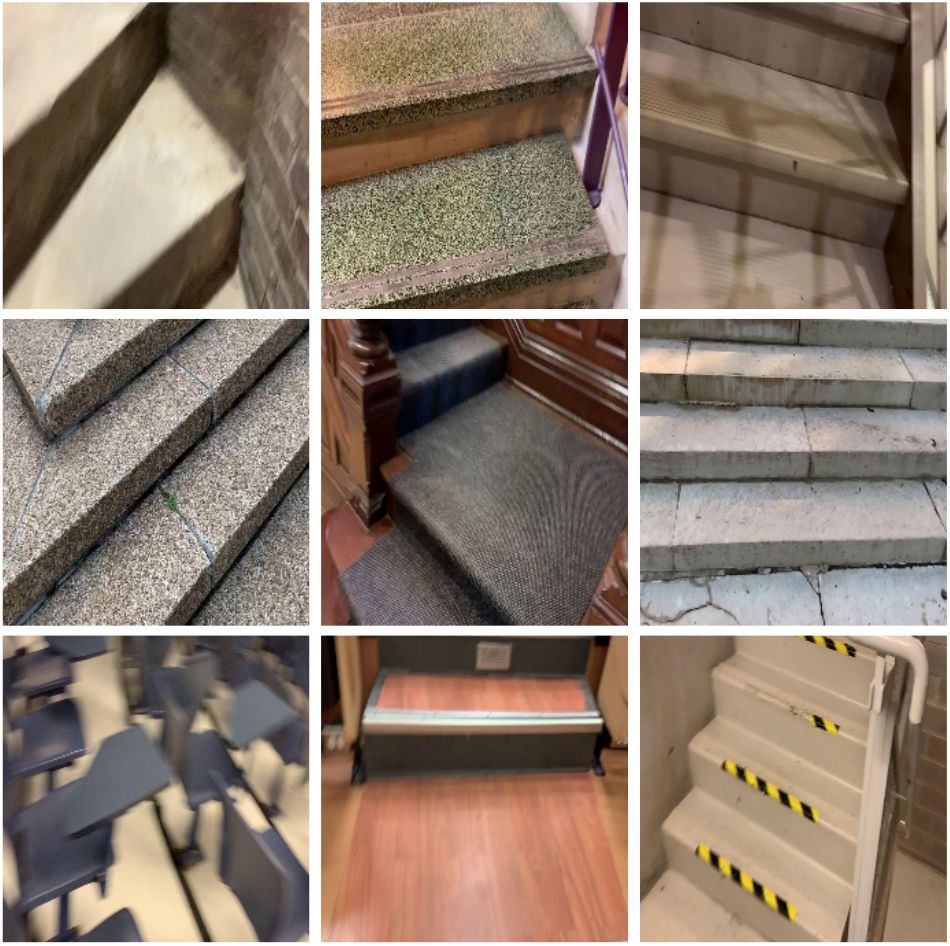
Examples of failure cases. The first row contains images of incline stairs (IS) that were misclassified as incline stairs – level ground (IS-LG). The second row contains images of incline stairs that were mis-classified as level ground -incline stairs (LG-IS). The third row contains different cases of level-ground misclassifications: the bottom-left image of level-ground was incorrectly classified as incline stairs and the bottom-middle and right images of level-ground—incline stairs were incorrectly classified as incline stairs.

## IV. Discussion

In this study, we developed a new perception system called *ExoNet-SSL* for human-robot walking using semi-supervised learning, which significantly reduced the number of required labelled training images while maintaining high prediction accuracy for stair recognition. Unlike previous studies that have been limited to supervised learning [7]-[15], we made use of manually labelled images from *StairNet* [8]-[9] as well as large amounts of unlabelled images (over 1.2 million images) from *ExoNet* [16] -the largest and most diverse open-source image dataset of real-world walking environments. Compared to supervised learning (98.4% accuracy) alike [8]-[9], our new semi-supervised learning model powered by mobile vision transformers achieved high classification accuracy during inference (98.8% accuracy) while requiring ∼35% less manually annotated images, thus improving overall training efficiency.

Despite these developments, our study has several limitations. While the number of labelled images is still relatively high, our work serves as a first step towards making computer vision more accessible to researchers in wearable robotics, especially on large-scale datasets. While a fully unsupervised model is a logical end-goal, semi-supervised learning can provide an initial milestone along this path. We also recognize the limitations of 2D RGB images and their lack of depth data that can be used for geometric feature extraction as the height of stairs can be important for adaptive control of robotic prosthetic legs and exoskeletons. Also, other algorithms besides FixMatch [19] could have been used to reduce the number of required labelled training images for semi-supervised learning. We could replace the pre-defined thresholds with flexible thresholds that can dynamically change (e.g., using curriculum pseudo labeling) to increase performance. Another limitation is that our vision transformer model, although lightweight and designed from efficient convolutional-transformer components, was not evaluated on an edge computing devices, as would be needed for real-time control of human-robot walk-ing. Future research should focus on improving model architecture, balancing accuracy across environmental states, and assessing the model inference speed on mobile and embedded devices. Unsupervised learning could eventually be used to improve performance and training efficiency. Methods such as invariant semantic information clustering [29] or cross-level discrimination for unsupervised feature learning could further decrease data labelling.

Another issue that we would like to explore in the future is balancing the image classification accuracies across different environment classes. Since we considered each of the four environmental states in our computer vision problem as equally important, we dedicated time to improving the model performance equaling across all classes. Although we significantly strengthened the prediction accuracies of minority classes using the focal loss and class weights, the overall image classification performance was inferior to the traditional, non-balanced approach. We describe this as the class accuracy allocation phenomenon and aim to study this behavior in future work.

In summary, the results of our new visual perception system powered by semi-supervised learning demonstrated the utility of leveraging large amounts of otherwise unused unlabelled data to improve training efficiency while maintaining high prediction accuracy for stair recognition. This research can help make computer vision and deep learning more accessible to researchers in wearable robotics and support the development of next-generation autonomous controllers for human-robot walking in real-world environments.

## Acknowledgment

We thank Nadiya Shvai from the National University of Kyiv-Mohyla Academy for sharing her expertise on semi-supervised learning for computer vision. This research is dedicated to the people of Ukraine in response to the 2022 Russian invasion and war.

## Notes

* Research supported by the Schroeder Foundation and the AGE-WELL Networks of Centres of Excellence (NCE) program, Canada.

https://ieee-dataport.org/documents/stairnet-computer-vision-dataset-stair-recognition

### Competing Interest Statement

The authors have declared no competing interest.

